# ShIVA – A user-friendly and interactive interface giving biologists control over their single-cell RNA-seq data

**DOI:** 10.1101/2022.09.20.508636

**Authors:** Rudy Aussel, Muhammad Asif, Sabrina Chenag, Sébastien Jaeger, Pierre Milpied, Lionel Spinelli

## Abstract

Single-cell technologies have revolutionised biological research and applications. As they continue to evolve with multi-omics and spatial resolution, analysing single-cell datasets is becoming increasingly complex. For biologists lacking expert data analysis resources, the problem is even more crucial, even for the simplest single-cell transcriptomics datasets.

We propose ShIVA, an interface for the analysis of single-cell RNA-seq and CITE-seq data specifically dedicated to biologists. Intuitive, iterative and documented by video tutorials, ShIVA allows biologists to follow a robust and reproducible analysis process, mostly based on the Seurat v4 R package, to fully explore and quantify their dataset, to produce useful figures and tables and to export their work to allow more complex analyses performed by experts.

## Introduction

In biology, single-cell RNA sequencing (scRNA-seq), and more generally single-cell genomics, have opened a new era. The scientific community has rapidly adopted those technologies as evidenced by their use in a growing number of high-impact biomedical publications (Svensson et al. 2020). The rapid growth of new applications (scRNA-seq, CITE-seq, scDNA-seq, spatial transcriptomics, scChIP-seq) is providing a large diversity of information about biological heterogeneity and dynamic processes in cell populations (Perkel 2021).

A major hindrance to the meaningful use of these technologies is that they require both a very good knowledge of the underlying biology and an expert level in data analysis (bioinformatics and mathematics). The successful analysis of single-cell genomics datasets is therefore the result of a close collaboration between biologists and bioinformaticians. This interdependence raises two main issues:

1. In early stages of the analysis, biologists need to rapidly browse the data to grasp a first idea of the underlying biology. To do so, they rely on various methods for data visualization, made available by bioinformaticians. Upon inspection of resulting reports, scientific experts often request refinements and new analyses to push further their understanding. During this iterative process, a significant amount of time is lost in the back-and-forth exchange between collaborators, and a first level analysis often requires several weeks.
2. ScRNA-seq data analysis challenges are quiet recent and many research groups are not fully equipped with required bioinformatics resources yet. In this context, in-depth analysis of experimental data is not always possible.

In recent years, the bottleneck between generation of experimental data and their analysis has been getting worse (Eisenstein et al. 2020). Until recently, the time required to produce datasets in the laboratory was greater than the time taken for bioinformatics analyses. Today, with the remarkable advances in biotechnologies, generating single-cell genomics datasets has become relatively easy and fast. On the other hand, time required for analysis has increased because of the large number and high complexity of analyses applied to those datasets. As a result, while research teams can produce large datasets quickly, they often have to wait long before analyzing them.

A tool enabling biologists to execute first steps of data analysis autonomously and deeply explore the biology of their datasets would considerably speed up the process, and allow them to improve the knowledge they can infer from the exploration. In this context, data analysis experts can focus on more complex analyses, to confirm biologists’ observations or bring new evidence. With that problem in mind, we have developed ShIVA, a Shiny Interface for Visualization and Analysis of single-cell datasets (Figure 1).

**Figure 1:**
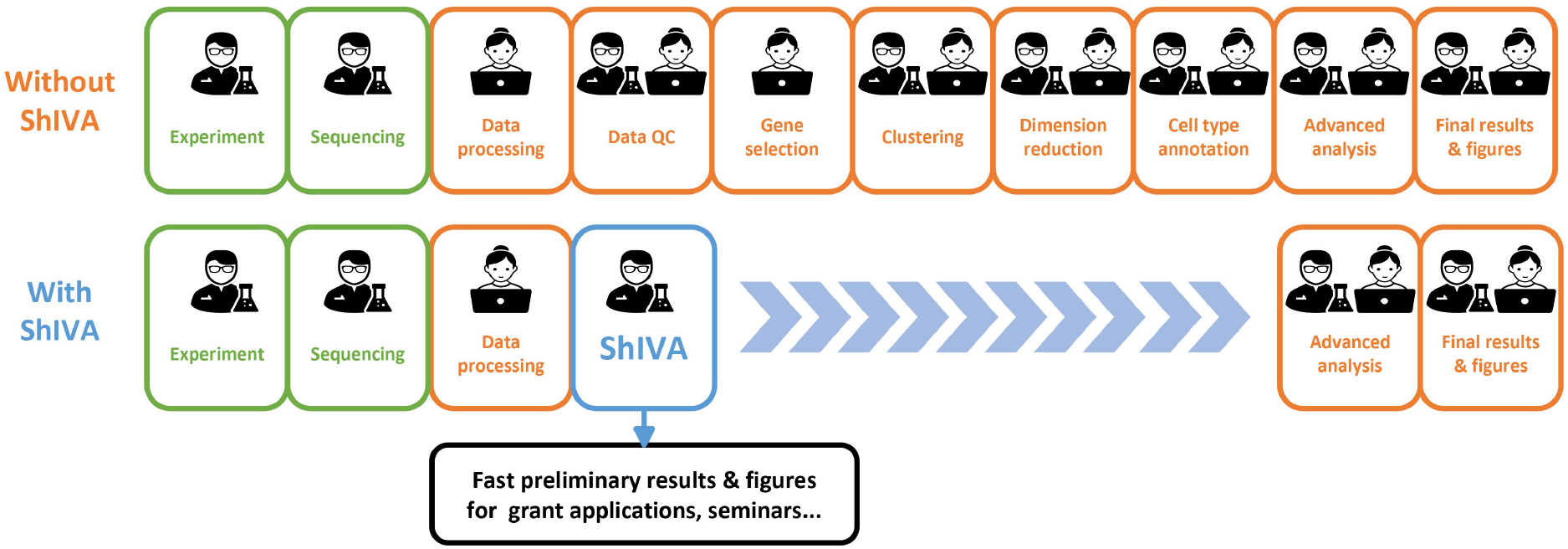
ShIVA reduces single-cell genomics data analysis time by placing the biologist project leader in power. ShIVA enables biologists to perform state-of-the-art exploratory analyses of their single-cell genomics datasets in a user-friendly interface.

## Results

### Principle of the interface

ShIVA is a software allowing any biologist to apply a correct and validated analytic process that corresponds to current state-of-the-art, relying mostly on functions from the Seurat v4 R package that was developed by single-cell data analysis experts (Hao et al., 2021). With this tool, non-experts (biologist researchers, PhD students or engineers) can load single-cell sequencing data (scRNA-seq or CITE-seq), apply the main quality control steps, execute normalization and dimensionality reduction, perform clustering, identify marker genes and their main functions, and finally visualize the data interactively to execute supervised and non-supervised scientific analyses.

ShIVA has been designed specifically to guide biologists through the analysis workflow, offering them to visualize, quantify and explore the data at each stage before making decisions affecting downstream analysis. The interface has been designed to be as user-friendly as possible while providing maximum choices. Specifically, ShIVA has been designed to allow the biologist to easily navigate through the workflow steps to revisit a decision, modify one or more parameters and iteratively refine their understanding of the data. Moreover, once the initial analysis steps have been completed, ShIVA offers a complete and fully customizable data exploration: the biologists can create the figures of their choice (maps, boxplots, violin plots, histograms, density plots, tables, with several options for style customization), quantify precisely what interests them most and create subsets of cells (for example a cell subtype) on which they can repeat the complete analysis process to deepen their understanding. The complete analysis can be saved as a “ShiVA project” file to be loaded later for further analysis.

ShIVA does not forget that collaboration with a data analysis expert will certainly be required for more advanced analyses. To ease this collaboration, ShIVA offers to export (i) all the figures, tables, list of genes or cells created by the biologists, (ii) a complete report of the analysis performed with all the details ensuring its reproducibility, and (iii) a file (R object) that can be re-used by the bioinformatician in an R environment to take up analyses where the biologist left them.

To help biologists use ShIVA correctly, we created tutorial videos for each step of the analysis workflow, available on a YouTube channel (https://www.youtube.com/channel/UCJJ3Svi8AY6XGx4Y9r9G3Iw).

### Overview of main features

ShIVA covers all the functions described in Figure 2. It is able to process data from scRNA-seq and CITE-seq experiments, including samples multiplexed with “hashtag” oligonucleotides (HTO). The workflow in ShIVA is performed through a succession of interdependent modules. Activating a module is allowed only if the mandatory previous steps have been correctly performed, helping the user to grasp the rigorous analysis process to follow. In each module, the user visualizes the relevant data, makes decision on the required analysis parameters and evaluates their impact through direct observation of the result. When the result of an analysis is not optimal, the user can easily step backward in the previous modules to modify the parameters and re-run the corresponding analysis. ShIVA keeps track of the user’s choice by defining a hierarchy of sub-projects, each of them containing the results of different user choices. Switching between sub-projects allows for comparison of analysis processes to optimize the deciphering of the dataset. Table 1 summarizes for each analysis module the parameters available to user customization, the different visualizations available, and the results.

**Figure 2:**
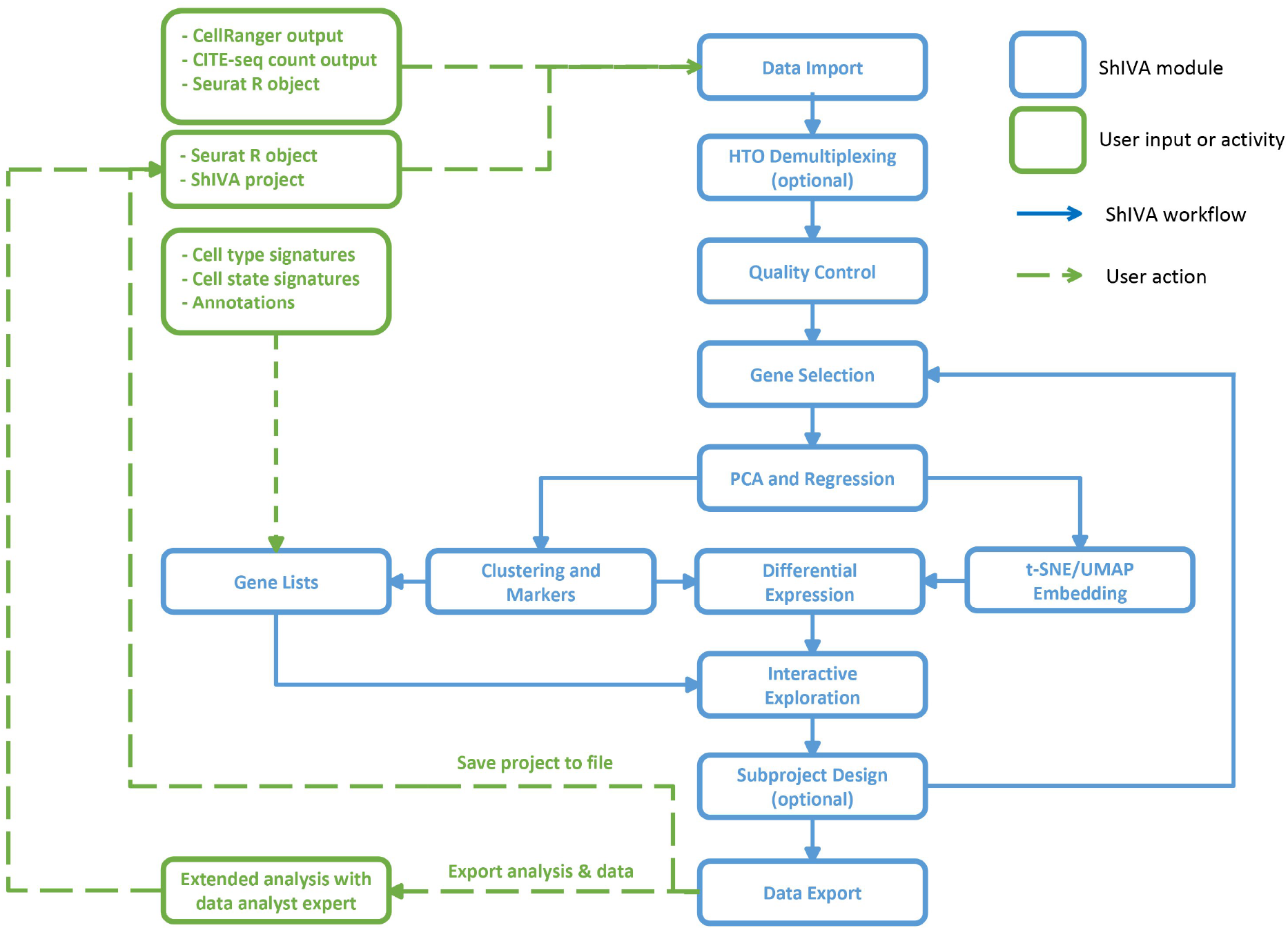
ShIVA modules and workflow. Schematic view of ShIVA analysis modules (blue boxes) enabling the analysis of single-cell genomics datasets from various standard user inputs (green boxes).

**Table 1:** List of ShIVA modules., Each row describes a module with its name (left column), the parameters available to user customization (middle column), and the different visualizations and results available (right column).

| **Module name** | **Parameters** | **Available visualizations / Results** |
| --- | --- | --- |
| **Data Import** | Project name, Species, Normalization method, Cells and genes thresholds | Creation of a ShIVA Project |
| **HTO demultiplexing** | Normalization method, quantile threshold (expert) | Number of cells by HTO, negative cells and multiplets; HTO UMAP plot; HTO Ridge plots |
| **Quality Control** | Range of : % of mitochondrial genes, % of ribosomal genes, Number of features per cell, Number of UMI per cell | Covariate violin plots; Covariate scatterplots; Number of cells filtered out |
| **Gene selection** | Gene selection method; Parameters of the selected method | Table of selected genes; scatterplot of selected genes |
| **PCA and Regression** | Number of PC to compute; Covariate to regress (optional) | Heatmap of loading genes per PC; Heatmap of PC and covariables correlations; Scatterplots of PC against covariables |
| **T-SNE/UMAP Embedding** | PCA analysis to use; Number of PC to use; Perplexity; Number of neighbors to consider | t-SNE plots; UMAP plots |
| **Clustering** | Number of PC to use; Clustering method; Parameters of the selected method; | Plots of the clusters in PCA or t-SNE or UMAP plots |
| **Markers genes** | Minimal expression percentage; Minimal log fold change; Analysis method | Table of all makers; Table of markers by cluster; Feature plots of top markers by cluster |
| **Differential expression** | Parameters to define the two group of cells; Minimal expression percentage; Minimal log fold change; Analysis method | Table of all DEG; Feature plots of top DEG; Functional enrichment of DEG in GO BP, MF, CC and KEGG pathways |
| **Interactive exploration - 2D plot** | Type of plot (PCA/t-SNE/UMAP); Variable to use for colors; Variables for layout (facet); Theme for axis and background; Size of dots; Selection of cells by manual gating (optional) | 2D plot with chosen colors, layout and theme; List of selected cells |
| **Interactive exploration - 1D plot** | Type of plots (violin/histogram/density); Variables to plot; Variable to use for colors; Theme for axis and background | 1D plot of the chosen variables, layout and theme |
| **Interactive exploration- table** | Type of table (counts/expression); Variables to compute in the table; Layout of table | Table of counts/expression of the chosen variables |

### Advantages of ShIVA

Several tools already exist to manage and visualize single-cell datasets (Jiang et al. 2022, Hasanaj et al. 2022, Moreno et al. 2021, BBrowser, CellxGene, SeqGeq …). To our knowledge, no freely available solution offers the combination of high level of simplicity and deep data exploration ShIVA provides. Existing user-friendly solutions are limited in the ability for user to control, compare and explore the analysis results. Compared to others tools, the main advantages of ShIVA are (i) user experience orientation, allowing a non-expert to easily understand and control each step of the analysis, (ii) modularity, allowing to easily follow the rigorous workflow of analysis, (iii) interactivity and iterability, allowing to acquire a better understanding of the underlying biology of the dataset, (iv) exportability and reproducibility, allowing to easily share and push forward the analysis with expert collaborators.

### Example of analysis with ShIVA

We tested ShIVA on a public dataset of human peripheral blood mononuclear cells (PBMC) from 8 donors (Figure 3A, as described in (Stoeckius et al., 2018)). Samples were stained with barcoded anti-CD45 hashtag antibodies prior to capture for 10x Genomics 3’v2 scRNA-seq. We downloaded gene expression and hashtag count matrices from the Seurat tutorial website (https://satijalab.org/seurat/articles/hashing_vignette.html) and combined both matrices into a Seurat object in R. The object was then loaded into ShIVA for analysis, with log normalization of the gene expression count matrix. We performed hashtag count matrix normalization and automatic demultiplexing to identify samples from the eight donors, and excluded doublets and negative events from further analysis (Figure 3B). We then performed gene expression quality controls, excluding cells expressing more than 5% mitochondrial gene transcripts and fewer than 100 genes (Figure 3C). We then selected 2,000 highly variable genes with the vst method, and performed principal component analysis (PCA) (Figure 3D). We selected the first 30 components for low dimension embedding and clustering. Cells from all donors were mixed in different clusters in the UMAP embedding (Figure 3E). After clustering with Leiden method at resolution 0.8, we discriminated 10 clusters corresponding to common peripheral blood cells that we annotated manually based on cluster-specific marker genes (Figure 3F). We visualized the number of genes detected per cell for all clusters as violin plots, highlighting that myeloid cells (clusters 1, 8 and 9) expressed more genes than lymphoid cells (Figure 3G). For higher resolution analysis of those myeloid cell clusters, we created a new object containing only cells from clusters 1, 8 and 9, and repeated the computation of 2,000 highly variable genes, PCA, UMAP embedding and clustering. That re-analysis revealed several sub-clusters of *CD14*+ monocytes, but did not reveal additional sub-clusters of *FCGR3A*+ monocytes or *CD1C*+ conventional dendritic cells (cDC) (Figure 3H). The exploratory analysis of that dataset in ShIVA took approximately 2 hours from loading the dataset to exporting all figure plots in vectorized pdf format.

**Figure 3:**
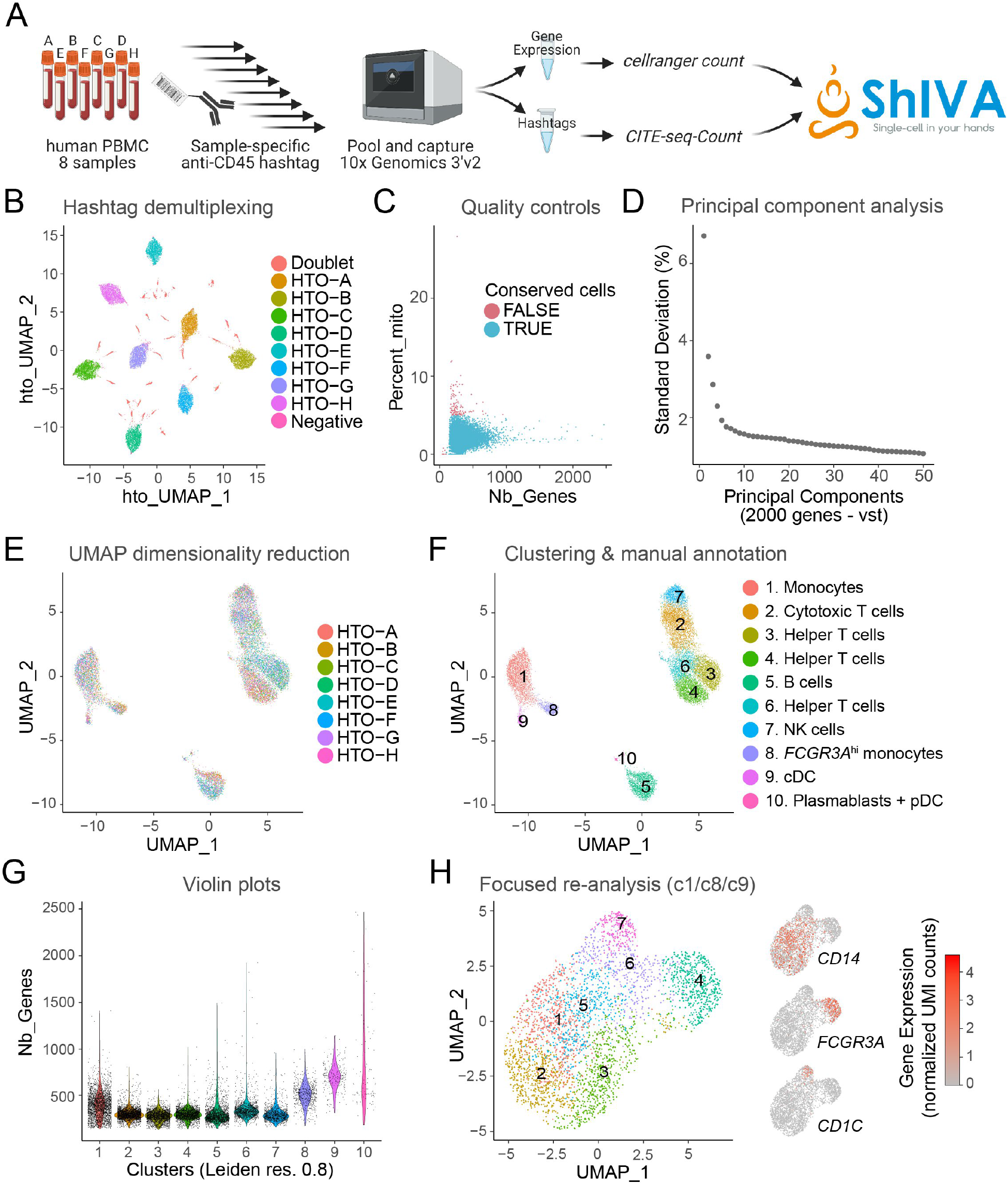
Example of analysis with ShIVA. **A**. Description of test dataset. Peripheral blood mononuclear cell (PBMC) samples from 8 human donors were stained with anti-CD45 hashtag antibodies, pooled and captured for 10x Genomics 3’v2 scRNA-seq and HTO library preparation, as described in (Stoeckius et al., 2018). Gene expression reads were processed with cellranger count to generate the cell x gene UMI count matrix, HTO reads were processed with CITE-seq-Count to generate the cell x HTO UMI count matrix. Both count matrices were downloaded from the Seurat tutorial website (https://satijalab.org/seurat/articles/hashing_vignette.html), combined into a Seurat object in R, and analyzed in ShIVA. **B**. UMAP computed on normalized HTO counts showing the results of HTO-based sample demultiplexing in ShIVA. Each dot is a cell, colored according to HTO-based sample assignation. Doublets and Negative cells are excluded from further analysis. **C**. Scatter plot of number of genes detected (x-axis) and percentage of transcripts from mitochondrial genes (y-axis) used for quality controls. Each dot is a cell, colored according to quality control results; only TRUE cells are conserved for further analysis. **D**. Scree plot of Standard Deviation percentage explained by Principal Components. Only the top 50 PC are shown, after computing PCA on the top 2000 highly variable genes defined by vst method. **E**. Gene expression based UMAP embedding computed on 30 PCs. Each dot is a cell, colored according to sample origin. **F**. Results of Leiden clustering presented on the gene expression based UMAP embedding. Each dot is a cell, colored according to cluster identity. Clusters were annotated manually based on cluster-specific marker genes. **G**. Violin plot of number of genes detected in cells according to cluster identity. **H**. Gene expression based UMAP embedding and clustering of myeloid cells after focused re-analysis of clusters 1, 8 and 9 from the analysis in **F**. The expression of discriminating marker genes *CD14, FCGR3A* and *CD1C* is shown as feature plots. Each dot is a cell, colored according to cluster identity (left) or gene expression levels (right).

## Discussion

In conclusion, ShIVA offers a user-friendly and user-oriented interface dedicated to the analysis of single-cell RNA-seq and single-cell CITE-seq data with an emphasis on ease of use, application of state-of-the-art methods, and reproducibility. ShIVA supports cell hashing analysis and provides great flexibility in visualization, whether by dimensionality reduction maps, boxplots, violin plots, histograms, density plots, or count tables. ShIVA is designed to give non-experts the power of state-of-the-art algorithms, allowing them to gain a deep understanding of their dataset while preparing for an optimized collaboration with data analyst experts in subsequent extensive analyses. A complete series of tutorial videos are available on YouTube (https://www.youtube.com/channel/UCJJ3Svi8AY6XGx4Y9r9G3Iw).

## Material and methods

ShIVA is a Shiny App (Beeley et al., 2018) built in an R framework (v. 4.0.3). ShIVA is deployed using Docker. ShIVA uses the Seurat package (v. 4.0.0, Hao et al. 2021) for the main analysis workflow. HTO demultiplexing is performed with MULTI-seq (McGinnis et al. 2019). Dimensionality reduction is performed using PCA (ade4 v.1.7.16, Dray et al. 2007), t-SNE (Van der Maaten et al. 2008) and UMAP (v. 0.2.7, McInnes et al. 2018). Clustering is performed through the methods proposed by Seurat among which the Leiden algorithm (v. 0.3.10, Traag et al., 2019). Visualization is performed using packages ggplot2 (v. 3.3.3), plotly (v.4.9.3), pheatmap (v 1.0.12) and heatmaply (v. 1.2.1). Complete details on the other R packages used are available in the dockerfile provided on GitHub.

## Acknowledgements

We thank Guillaume Voisinne for his help in designing the basic architecture of the software. We thank Bernard Malissen for supporting the project through human resources. We would like to thank all our beta-testers from Centre d’Immunologie de Marseille-Luminy, Institut de Neurobiologie de la Méditérranée, Institut de Biologie du Développement de Marseille, Centre de Recherche en Cancérologie de Marseille, Centre de Recherche en CardioVasculaire et Nutrition, Theories and Approaches of Genomic Complexity, Unité de biologie Moléculaire, Cellulaire et du Développement, Institut de Recherche Saint Louis, Institut de Biologie Paris-Seine, RESTORE, and Institut de la Vision.

## Funding

The ShIVA project has been funded by the Union’s Horizon 2020 Framework Programme (grant agreement 787300 [BASILIC] to B. Malissen), the CENTURI Tech Transfer program and the Avesian ITMO GGB Amorçage program.

## Code availability and tutorials

Code and documentation are available from the GitHub repository: https://github.com/CIML-bioinformatic/ShIVA.

Complete video tutorials are available on YouTube: https://www.youtube.com/channel/UCJJ3Svi8AY6XGx4Y9r9G3Iw

Docker image is available on DockerHub: https://hub.docker.com/repository/docker/cb2m/shiva

## Author contributions

SJ, PM and LS designed the project, produced the feature specifications and supervised the development. RA, MA and SC developed the software. PM and LS wrote the manuscript with input from all authors. All authors read and approved the manuscript.

## Notes

### Competing Interest Statement

The authors have declared no competing interest.

## References

BBrowser : https://bioturing.com/bbrowser

Beeley, C., & Sukhdeve, S. R. (2018). Web Application Development with R Using Shiny: Build stunning graphics and interactive data visualizations to deliver cutting-edge analytics. Packt Publishing Ltd.

CellxGene : https://github.com/chanzuckerberg/cellxgene

Dray, Stéphane, and Anne-Béatrice Dufour. “The Ade4 Package: Implementing the Duality Diagram for Ecologists.” Journal of Statistical Software, vol. 22, no. 4, 2007, https://doi.org/10.18637/jss.v022.i04.

Eisenstein, Michael. “Single-Cell RNA-Seq Analysis Software Providers Scramble to Offer Solutions.” Nature Biotechnology, vol. 38, no. 3, Mar. 2020, pp. 254–57, https://doi.org/10.1038/s41587-020-0449-8.

Hao, Yuhan, et al. “Integrated Analysis of Multimodal Single-Cell Data.” Cell, vol. 184, no. 13, June 2021, pp. 3573-3587.e29, https://doi.org/10.1016/j.cell.2021.04.048.

Hasanaj, Euxhen, et al. “Interactive Single-Cell Data Analysis Using Cellar.” Nature Communications, vol. 13, no. 1, Apr. 2022, p. 1998, https://doi.org/10.1038/s41467-022-29744-0.

Jiang, Andrew, et al. “ICARUS, an Interactive Web Server for Single Cell RNA-Seq Analysis.” Nucleic Acids Research, May 2022, p. gkac322, https://doi.org/10.1093/nar/gkac322.

McGinnis, Christopher S., et al. “MULTI-Seq: Sample Multiplexing for Single-Cell RNA Sequencing Using Lipid-Tagged Indices.” Nature Methods, vol. 16, no. 7, July 2019, pp. 619–26, https://doi.org/10.1038/s41592-019-0433-8.

McInnes, Leland, et al. UMAP: Uniform Manifold Approximation and Projection for Dimension Reduction. 2018, https://doi.org/10.48550/ARXIV.1802.03426.

Moreno, Pablo, et al. “User-Friendly, Scalable Tools and Workflows for Single-Cell RNA-Seq Analysis.” Nature Methods, vol. 18, no. 4, Apr. 2021, pp. 327–28, https://doi.org/10.1038/s41592-021-01102-w.

Perkel, Jeffrey M. “Single-Cell Analysis Enters the Multiomics Age.” Nature, vol. 595, no. 7868, July 2021, pp. 614–16, https://doi.org/10.1038/d41586-021-01994-w.

SeqGeq : https://www.flowjo.com/solutions/seqgeq

Stoeckius, Marlon, et al. “Cell Hashing with Barcoded Antibodies Enables Multiplexing and Doublet Detection for Single Cell Genomics.” Genome Biology, vol. 19, no. 1, Dec. 2018, p. 224, https://doi.org/10.1186/s13059-018-1603-1.

Svensson, Valentine, et al. “A Curated Database Reveals Trends in Single-Cell Transcriptomics.” Database: The Journal of Biological Databases and Curation, vol. 2020, Nov. 2020, p. baaa073, https://doi.org/10.1093/database/baaa073.

Traag, V. A., et al. “From Louvain to Leiden: Guaranteeing Well-Connected Communities.” Scientific Reports, vol. 9, no. 1, Mar. 2019, p. 5233, https://doi.org/10.1038/s41598-019-41695-z.

Van der Maaten, Laurens, Hinton, Geoffrey, Visualizing data using t-SNE, Journal of Machine Learning Research, 9(2605):2579–2605, 2008.

